# Direct immunosensing of avian influenza A virus in whole blood using hybrid nanocomposites

**DOI:** 10.1101/343376

**Authors:** John Buozis, Syed Rahin Ahmed, Rohit Chand, Éva Nagy, Suresh Neethirajan

## Abstract

A sandwich-based electrochemical immunosensor was designed for detection of avian influenza virus (AIV) strains H5N1 and H4N6. This sensor was developed using gold-graphene nanocomposites, immobilized viral antibodies, and CdTe quantum dot electrochemical tagging. The nanocomposites were formed by the simultaneous reduction of a gold salt and graphene using hydroquinone as the reducing agent, thus producing non-spherical gold nanoparticles on graphene sheets. Viral antibodies were immobilized on nanocomposites and CdTe quantum dots through N-(3-dimethylaminopropyl)-N′-ethylcarbodiimide and N-hydroxysuccinimide chemistry. Cyclic voltammetry studies were used to validate the detection of H5N1 surface protein and H4N6 inactivated virus. The immunosensor detected H5 protein in phosphate buffer solution (pH 7.4) with a limit of detection (LOD) of 10 fg/mL and a linear detection range was established for 10 ng/mL to 10 pg/mL. The biosensor detected H4N6 in three parts diluted whole chicken blood with a LOD of 1.28×10^−7^ hemagglutinating units (HAU). Commercial ELISA testing for H5N1 and H4N6 showed limits of detection of 10 ng/mL and 0.128 HAU, respectively. The sensor showed 10^6^-fold increased detection of H4N6 virus in blood in comparison to its commercial ELISA kit counterpart. The developed immunosensor effectively change the way avian influenza is detected, monitored, and controlled; transforming time-consuming reactive methods, into rapid predictive technology.

## 1. Introduction

To meet the growing demands for animal protein, global poultry production will double in the next 25 years. The global poultry industry has been deeply impacted by outbreaks of avian influenza virus (AIV) since the late 1990’s. The Canadian poultry industry has also had its share of major losses due to AIV, most notably was the 2004 outbreak in British Columbia, which resulted in culling 19 million birds. Recent AIV outbreaks in British Columbia and Ontario in 2015 also have caused economic losses to the Canadian poultry industry. Aside from the significant impact of AIV on animal health, some of these viruses have an impact on public health. AIV causes three- to five-million people to fall severely ill, resulting in 250,000 to 500,000 fatal cases annually in developing countries (World Health Organization, 2017a). Furthermore, hemagglutinin (HA) and neuraminidase (NA) surface protein combinations are used to characterize influenza viruses. There are 18 HA (H1 – H18) and 11 NA (N1 – N11) subtypes, respectively. Of these subtypes, H5 and H7 are of major concern within the scientific community; as they manifest as low pathogenic infections in waterfowl, which can become highly pathogenic when introduced to domestic poultry, and are zoonotic. (Canadian Food Inspection Agency, 2015; Centers for Disease Control and Prevention, 2017, 2015; Health canada, 2008; Jensen et al., 2013; Olsen et al., 2006; World Health Organization, 2017b; Zhu et al., 2014). Consequently, governments and farmers alike are under immense pressure to ensure the health of poultry and poultry consumers.

Preventing the spread of avian influenza infection is the best way to keep disease outbreaks under control. Prevention starts with effective bio-surveillance through early disease diagnosis. To date there are no pen-side or coop-side tests available for rapid diagnosis. Conventional methods of avian influenza detection include one-step reverse transcription polymerase chain reaction (RT-PCR), hemagglutinin inhibition tests, enzyme-linked immunosorbent assay (ELISA), embryonated egg virus culturing, and chicken pathogenicity tests (Jensen et al., 2013; United States Department of Agriculture, 2015; World Organization for Animal Health, 2016).

In the United States, tests carried out by the National Animal Health Laboratory Network (NAHLN) (United States Department of Agriculture, 2015) are as follows: matrix screening for AI viruses, H5 subtype screening, H7 subtype screening, and N1 subtype screening; all of which are RT-PCR based tests (American Plant Health Inspection Service, 2008; United States Department of Agriculture, 2015). These tests are followed by three types of confirmatory tests: virus isolation tests (in embryonated eggs), genetic sequencing tests, and chicken pathogenicity tests. The test samples are usually obtained from fecal or tracheal swabs from live specimens. Typically these tests take 2-3 weeks to run, require expensive equipment, and require highly trained technicians (American Plant Health Inspection Service, 2008).

To overcome the obstacles of poor diagnostic turnaround and the need for specialized facilities, research has been moving towards virus detection on the nanoscale using point-of-care (POC) biosensors (Neethirajan et al., 2017). The major benefits of nanoscale virus detection include: a significant reduction in reaction time due to increased surface area for the reaction to take place; and a significant reduction in the costs of testing (e.g. reagent costs, personnel costs, facility costs, and transportation costs). Due to the current technology limitations, this work will focus on bridging the gap through the design of a rapid point-of-care biosensor for the detection of avian influenza A viruses.

Graphene is an abundant, inexpensive two-dimensional atomic crystal with outstanding physical properties, including extreme mechanical strength, exceptionally high electronic conductivities, superior surface area, and biocompatibility. It is an excellent substrate for biomolecule anchoring and detection due to its surface area of 2630 m2/g10 and unique sp2 (sp2/sp3) bonded network (Hu et al., 2015;Veerapandian and Neethirajan, 2015). In addition, by exploiting the electrochemical properties, graphene can be functionalized easily for developing novel biosensing and transduction mechanisms. Recent graphene-based biosensing platforms developed in our lab and others (Veerapandian et al., 2016a, 2016b; Weng and Neethirajan, 2016) indicate the potential for an electrochemical nanobiosensing platform for virus detection applications.

In this study, an immunosensor was designed by incorporating gold-graphene nanocomposites, antibody-antigen immunochemistry, and electrochemical quantum dot tagging. The goal of this study was to develop a sensing mechanism that was more sensitive and less time consuming than commercial ELISA. The proposed immunosensor uses a thin film fabricated of gold-graphene nanocomposites on a screen-printed electrode. Virus-specific antibodies were immobilized on the nanocomposite surface using N-(3-dimethylaminopropyl)- N′-ethylcarbodiimide (EDC) and N-hydroxysuccinimide (NHS) carbodiimide chemistry. Cadmium telluride (CdTe) quantum dots were conjugated with virus-specific antibodies in an *in-situ* manner using EDC/NHS chemistry. This sensing mechanism is an immunosensing on a screen-printed electrode, in which the magnitude of the CdTe electrochemical signal is proportional to the antigen concentration. The immunosensor was first designed for H5N1 viral protein as a proof-of-concept. To demonstrate the practicality of the immunosensor for real virus detection, low pathogenic H4N6 spiked in whole chicken blood was studied.

## 2. Materials and methods

### 2.1 Materials and reagents

Gold (III) chloride trihydride (HAuCl4·3H2O), L-polylysine, hydroquinone, potassium hexacyanoferrate (K_4_[Fe(CN)_6_]), potassium hexacyanoferrite (K_3_[Fe(CN)_6_]), phosphate buffer saline (PBS), gold cleaning solution, N-(3-dimethylaminopropyl)-N′-ethylcarbodiimide (EDC), N-hydroxysuccinimide (NHS), CdTe quantum dots and pyrene carboxylic acid were purchased from Sigma-Aldrich (MO, USA). Graphene (4% wt. water dispersion) was obtained from ACS Materials (CA, USA). All the chemicals were of analytical grade and used as received without further purification. Screen-printed gold electrodes were purchased from Dropsens (Spain). Whole chicken blood and influenza virus A (H1N1) surface protein were purchased from Cedarlane Labs (ON, Canada). Anti-influenza A (H5N1) virus hemagglutinin (HA) antibody and influenza virus A (H5N1) surface protein were purchased from Abcam, Inc., (Cambridge, UK). Anti-H4 (H4N6) polyclonal antibody was purchased from MyBioSource Inc., (San Diego, USA). Milli-Q water (18.2 MΩ, DI water) was used throughout the experiments.

### 2.2 Avian influenza (H4N6) and (H9N2) virus cultures

Low pathogenic AIV H4N6 (avian influenza A/Duck/Czech/56 (H4N6)) was propagated in 11-day-old embryonated chicken eggs by inoculation into the allantoic cavity (Szretter et al., 2006). Infectious titer in allantoic fluid was determined at 72 h post-inoculation and expressed as a 50% tissue culture infective dose 128 HAU/50 µL.

Inactivated AIV H9N2 (A/Turkey/Ontario/1/66) was propagated in 11-day-old embryonated SPF chicken eggs. The egg-derived virus was inactivated with formalin (final concentration 0.02%) for 72 h at 37 °C. The protein content of the inactivated virus preparation was determined using haemagglutination inhubition (HI) assay and expressed as 50% tissue culture infective dose 128 HAU/50 µL (Singh et al., 2016).

### 2.3 Fabrication of non-spherical graphene-gold nanocomposites

The nanocomposites were synthesized in a one-pot, *in-situ* method resulting in a final working solution volume of 20 mL. First, 18 mL of 40X diluted graphene solution (final solution concentration of 1 mg/mL) was sonicated for 15 minutes to separate the graphene sheets. Next, 1 mL of HAuCl_4_ (final solution concentration of 2.5 × 10^−4^ M) was added to the graphene solution under constant stirring. The solution was then stirred for 30 minutes. Next, 1 mL of hydroquinone (final solution concentration of 2.5 × 10^−4^ M) was added to the graphene gold solution to simultaneously reduce Au^3+^ to Au^0^ and graphene to reduced graphene. The solution was stirred for 1 hour at room temperature to allow complete reduction. The solution was then centrifuged at 15,000 rpm for 5 minutes to remove any unused reactants. The supernatant was removed from each tube, followed by a washing step with DI water. This centrifugation and washing procedure was carried out three times to ensure that any remaining reducing agent had been removed. The solution was then returned to a single 20 mL glass vile after the final washing step. The nanocomposite solution was then stored in a refrigerator at 4°C for future use.

The graphene-Au nanocomposites were characterized using UV-Visible spectroscopy (Cary 100, Agilent Technologies), transmission electron microscopy (TEM, FEI Tecnai G2 F20 microscope), scanning electron microscopy (SEM), energy dispersive x-ray (EDX) analysis, and electrochemical technique.

### 2.4 Nanocomposite deposition on electrodes

All electrodes were first cleaned by dropping 10 µL of gold cleaning solution onto the working electrode. After 10 seconds the electrodes were washed thoroughly with DI water. Next, 5 µL of L-polylysine was dropped onto the working electrode and was spread to cover the entire working electrode area. The electrodes were covered in a petri dish dried for 2 hours at room temperature, followed by a DI water rinse to remove any unbound L-polylysine. The same process was carried out for depositing the graphene-Au nanocomposite solution. These alternating layers formed L-polylysine/nanocomposite bilayers on the substrate. Two bilayers were used (meaning 4 layers in total) for the electrochemical immunosensing.

### 2.5 Antibody conjugation

Electrodes were first functionalized by using 1 mM pyrene carboxylic acid (adding a - COOH group to the graphene sheets). 10 µL of pyrene carboxylic acid was deposited onto each working electrode and the electrodes were set to dry for 1 hour. Next, 5 µL of 4 mM EDC was deposited onto the working electrodes followed by 5 µL of 10 mM NHS. The electrodes were left for 10 minutes to allow reaction between EDC and NHS, which was followed by a light DI water wash step to remove any o-acylisourea by-product. Next, 5 µL of the respective primary antibody, anti-H_x_ (1 µg/mL), was deposited onto the working electrode. The electrodes were incubated overnight at 4°C in a moisture chamber.

CdTe quantum dots were conjugated with anti-N1 antibodies (1 µg/mL) using the same EDC/NHS carbodiimide crosslinking. The quantum dots were also incubated overnight at 4°C. The same procedures were followed to immobilize Anti-H4 antibodies onto the CdTe quantum dots.

### 2.6 Optimization of AIV immunosensor

The concentration of primary antibodies was tuned by conducting cyclic voltammetry (CV) studies on electrodes with varying antibody concentrations (0.5 µg/mL to 2.5 µg/mL). A 1:1 mixture of 5 mM of K_4_[Fe(CN)_6_] and K_3_[Fe(CN)_6_] in 1X PBS (pH 7.4) was used as the electrolyte solution. During testing 100 µL of the electrolyte solution was dropped onto the electrode. Antibody concentration that resulted in the lowest current corresponding to highest surface coverage was chosen for the design. A commercial potentiostat (DRP-STAT200, Dropsens, Spain) was used to measure electrochemical redox current at a scan rate of 0.01 V/s. Similarly, a study was conducted using various antibody-antigen interaction times (0 min, 5 min, 10 min, 20 min, and 60 min). For this test 5 µL of target antigen (H4N6) spiked blood dilution was dropped onto each respective electrode, immediately followed by 5 µL of CdTe anti-H4 bioconjugate. Incubation time was considered to begin after the CdTe bioconjugates were added. After each incubation time period, the electrodes were gently washed with DI water. Each electrode was tested by micropipeting 100 µL of PBS buffer onto the antibodyantigen-antibody superstructure. The electrochemical redox current of the CdTe quantum dot reporters was measured to obtained applicable incubation time.

### 2.7 Electrochemical immunosensing of AIV

The schematic of electrochemical immunosensing of AIV is illustrated in Fig. 1. Screen-printed gold electrodes were used for these tests. A total of 8 serial dilutions of H5N1 surface protein were used (1 µg/mL to 1 fg/mL), each concentration was tested in triplicates. Similarly, a total of 8 serial dilutions of H4N6 were used (128 HAU, 1.28 HAU, 1.28 × 10^−2^ HAU, 1.28 × 10^−3^ HAU, 1.28 × 10^−4^ HAU, 1.28 × 10^−5^ HAU, 1.28 × 10^−6^ HAU, and 1.28 × 10^−7^ HAU). H4N6 dilutions were spiked in three-parts diluted whole chicken blood (PBS, pH 7.4). 5 µL of each viral dilution was dropped onto respective anti-H_x_ modified working electrode. Next, 5 µL of the anti-N1 (for H5N1) or anti-H4 (for H4N6) conjugated CdTe quantum dots was micropipetted onto each working electrode. In case of H5N1, the electrodes were incubated in a moisture chamber for 1 hr at 4°C to allow for antibody-antigen interaction, which was then followed by a DI water wash to remove any unbound quantum dot reporters. The H4N6 virus was incubated for an optimized duration of 10 minutes. The electrochemical redox current of the CdTe quantum dot reporters was measured with varying antigen concentrations. Each electrode was tested by micropipeting 100 µL of PBS buffer onto the antibody-antigen-antibody superstructure. All tests were conducted using a scan range of 0.1 V to -1 V with a scan rate of 0.01 V/s and a sampling rate of 0.002 V/s.

**Fig. 1.**
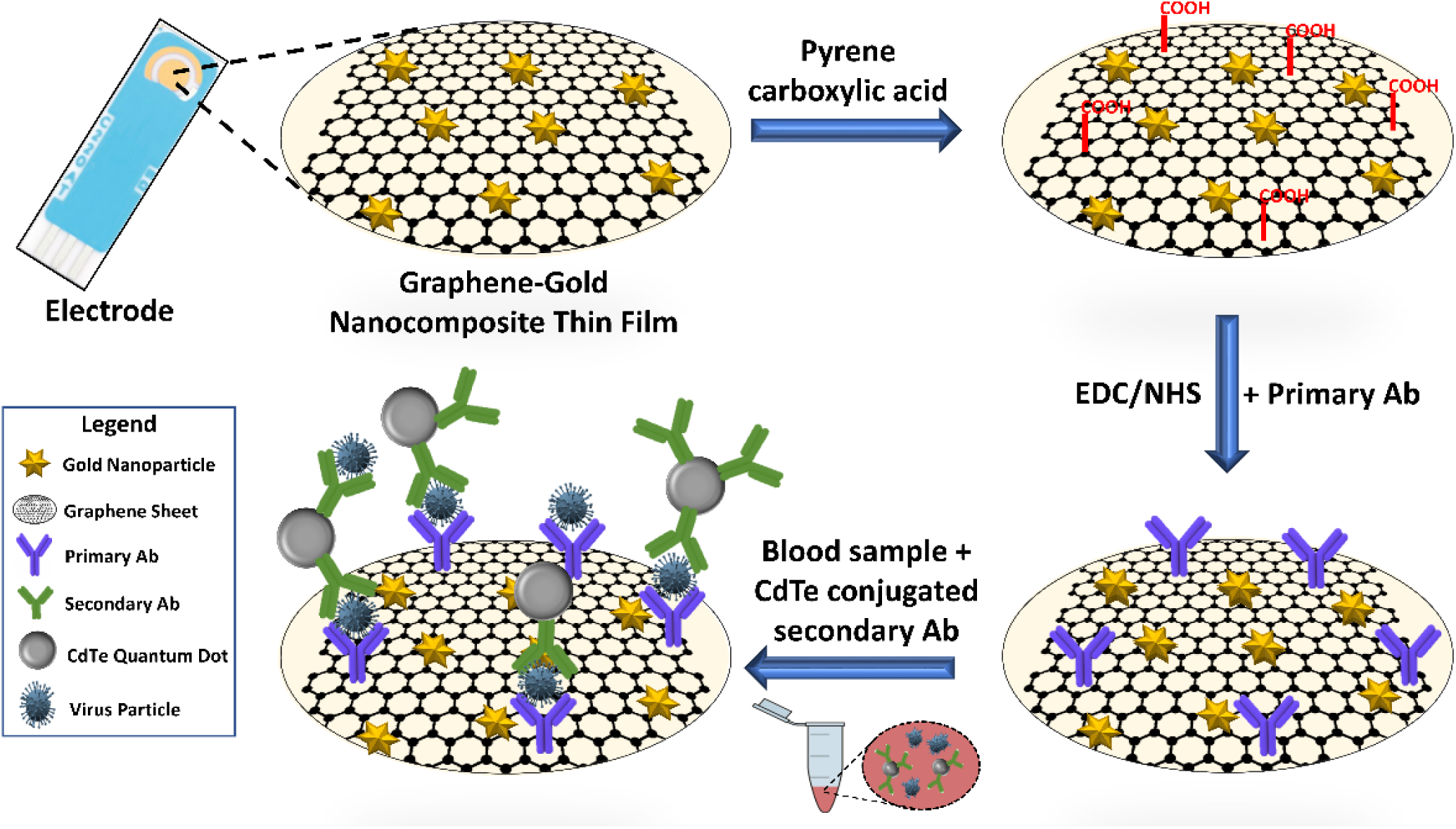
Electrochemical immunosensing mechanism for direct detection of influenza A virus in a whole chicken blood sample matrix.

### 2.8 Specificity and cross reactivity studies

The designed immunosensor was tested for specificity to H5N1 and H4N6. This was done by using H1N1 recombinant protein, H9N2 virus, and 1 mg/mL peptidoglycan (dispersion in 1X PBS) as negative controls. H5N1 recombinant protein was also used as a negative control for H4N6 virus. The control concentrations used for recombinant proteins and viruses were 1 µg/mL and 128 HAU, respectively. For cross reactivity testing of H4N6 in diluted whole blood, the sample was spiked with 1 mg/mL peptidoglycan and 128 HAU of H4N6. Blanks were also run for both H5N1 and H4N6 as negative controls. These tests were conducted in triplicates.

### 2.9 Validation studies with commercial ELISA

A comparison study was performed with a commercial avian influenza A H4N6 (Cat. No: NS-E10156, Novatein Biosciences, Woburn, MA, USA) ELISA Kit to validate the designed immunosensor. Various virus titers were prepared using sample diluent provided in the ELISA kit box and by strictly following the manufacturer’s protocol in the performance of the bioassay.

## 3. Results

### 3.1 Nanocomposite characterization

In this study, graphene sheets were decorated with non-spherical nanoparticles in a one-pot *in-situ* simultaneous reduction of graphene and a gold salt using hydroquinone as a reducing agent. TEM images of the fabricated nanostructures can be observed in Fig. 2. From Fig. 2, more particularly panels C and D, it can be confirmed that graphene sheets have been decorated with non-spherical or “spikey” nanoparticles. The resulting nanocomposites were also examined using SEM and EDX to determine their composition (Fig. S1). Through elemental analysis, it was found that C, Au, and O were present in the sample, indicating that the graphene sheets were successfully decorated with gold nanoparticles. This was also confirmed via UV-visible spectroscopic studies (Fig. S2). From Fig. S2, the nanocomposite exhibited the characteristic π – π bond of the polyaromatic C – C at 230 nm for graphene, as well as a second broad peak that was associated with the gold nanoparticles. It is hypothesized that using hydroquinone as a reducing agent would promote the formation of non-spherical gold nanoparticles on graphene sheets. The reason non-spherical gold nanoparticles were desired was because they would act as excellent spacers between the graphene sheets, thus reducing π – π stacking, promoting inter-layer linkage, and increasing the conductivity of graphene.

**Fig. 2.**
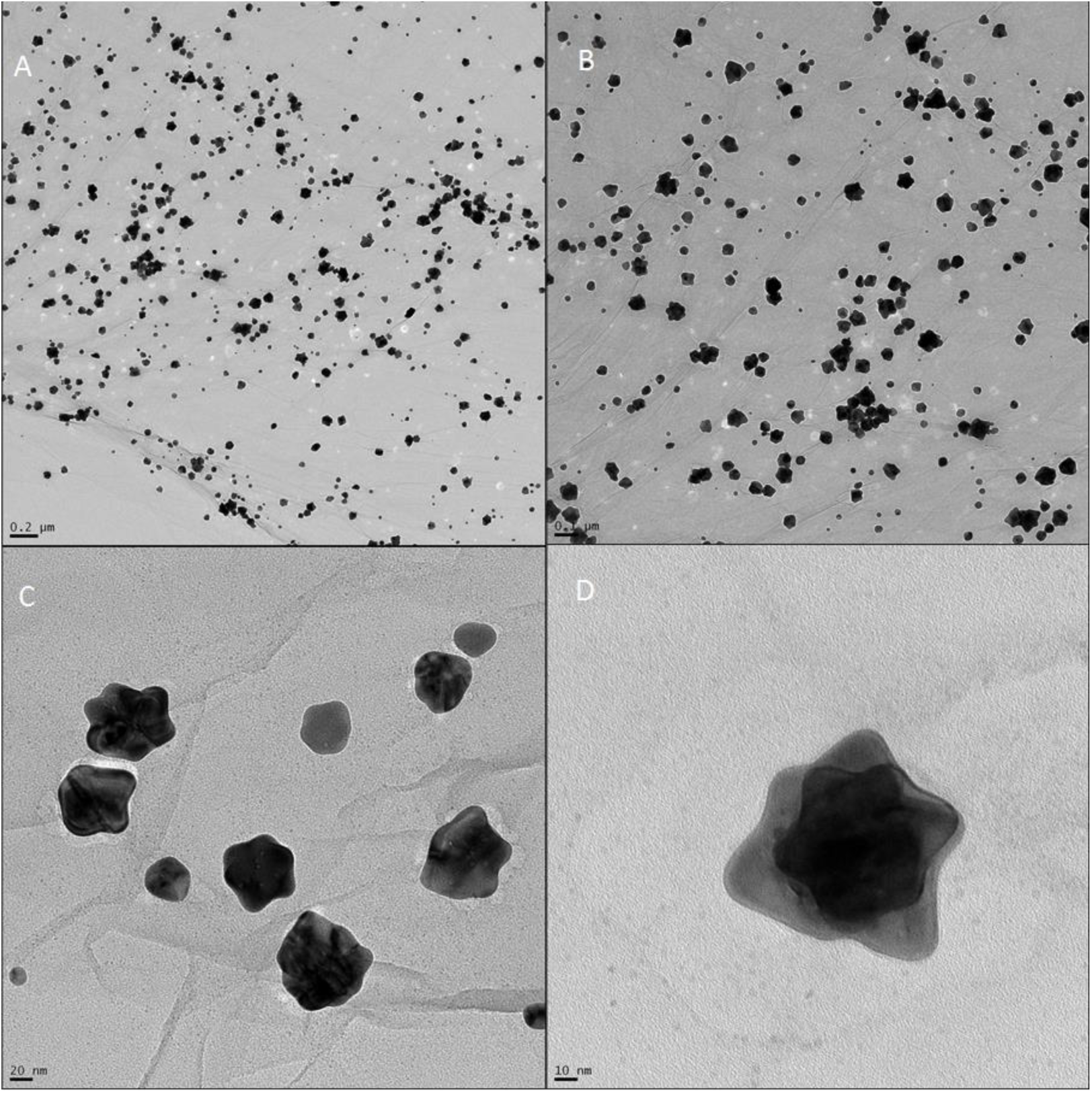
TEM images of non-spherical graphene-gold nanocomposites. (A) 200 nm scale demonstrates that graphene has been decorated with numerous nanoparticles; (B) 100 nm scale demonstrates non-spherical shape; (C) 20 nm scale depicts a small group of non-spherical nanoparticles as well as detailed graphene sheets; and (D) 10 nm scale depicts a single non-spherical nanoparticle.

Cyclic voltammetry (CV) studies were conducted to determine the signal amplification due to electrode surface modifications with the developed nanocomposite (Fig. S3). From Fig. S3, it can be observed that each successive bilayer (L-polylysine and nanocomposite) caused an associated response. We found that a single bilayer would sometimes wash away after testing, therefore additional bilayers were needed. The second and third bilayers provided more stable surface modifications while also increasing the measured signal. However, the two bilayer modification was chosen for further experimentation, as it reduced electrode fabrication time, provided a stable modification, and increased the measured signal.

### 3.2 Antibody concentration optimization

Five different antibody concentrations were used to determine an optimum coverage of the working electrode by the primary antibody. The concentration studies can be observed in Fig. S4. The primary antibody concentration with the lowest current value was found to be 1 µg/mL. This was chosen as the standard as experimentation was continued. The same concentration of secondary antibody was used to form CdTe quantum dot bioconjugates.

The proposed immunosensor must provide sensing results more rapidly than current conventional methods. In attempt to reduce the time for analysis, antigen – antibody interaction time studies were conducted (Fig. S5). It was found that the highest response is received at an incubation period of 60 minutes. This is on-par with current conventional techniques, however, we wanted a sensor that could be used rapidly in a point-of-care fashion. Therefore, we chose and incubation time of 10 minutes, as it provided the second highest response. An incubation time of 10 minutes is much more feasible than 60 minutes when it comes to point of care device employment. Therefore, an incubation time of 10 minutes was used for the H4N6 study.

### 3.3 H5N1 protein detection in 1X PBS

Detection of H5N1 recombinant protein was used as a proof of concept to determine if the immunosensor mechanism would work. The mechanism was later adapted to detect H4N6 virus. The CV profiles of the various H5N1 protein concentrations are shown in Fig. 3(A). Upon conducting H5N1 sensing experiments, two negative characteristic peaks were found in the CV profiles, one at -0.35V and the second at -0.75V, corresponding to the CdTe bioconjugate reporters. It was found that the characteristic peaks at -0.75V were more prominent, and thus were used to obtain the current – antigen concentration data shown in Fig. 3(B). Fig. 3(B) shows a near sigmoidal relationship between peak current and recombinant protein concentration. It was found that the biosensor could distinctly detect spiked concentrations of H5N1 protein in 1X PBS (pH 7.4) from 1 µg/mL to 10 fg/mL. A linear range exists between 10 ng/mL and 10 pg/mL of recombinant protein (Fig. 3(B) inset). The coefficient of determination and slope values for this relationship were found to be 0.987 and 0.9242 µA*mL*µg^−1^, respectively.

**Fig. 3:**
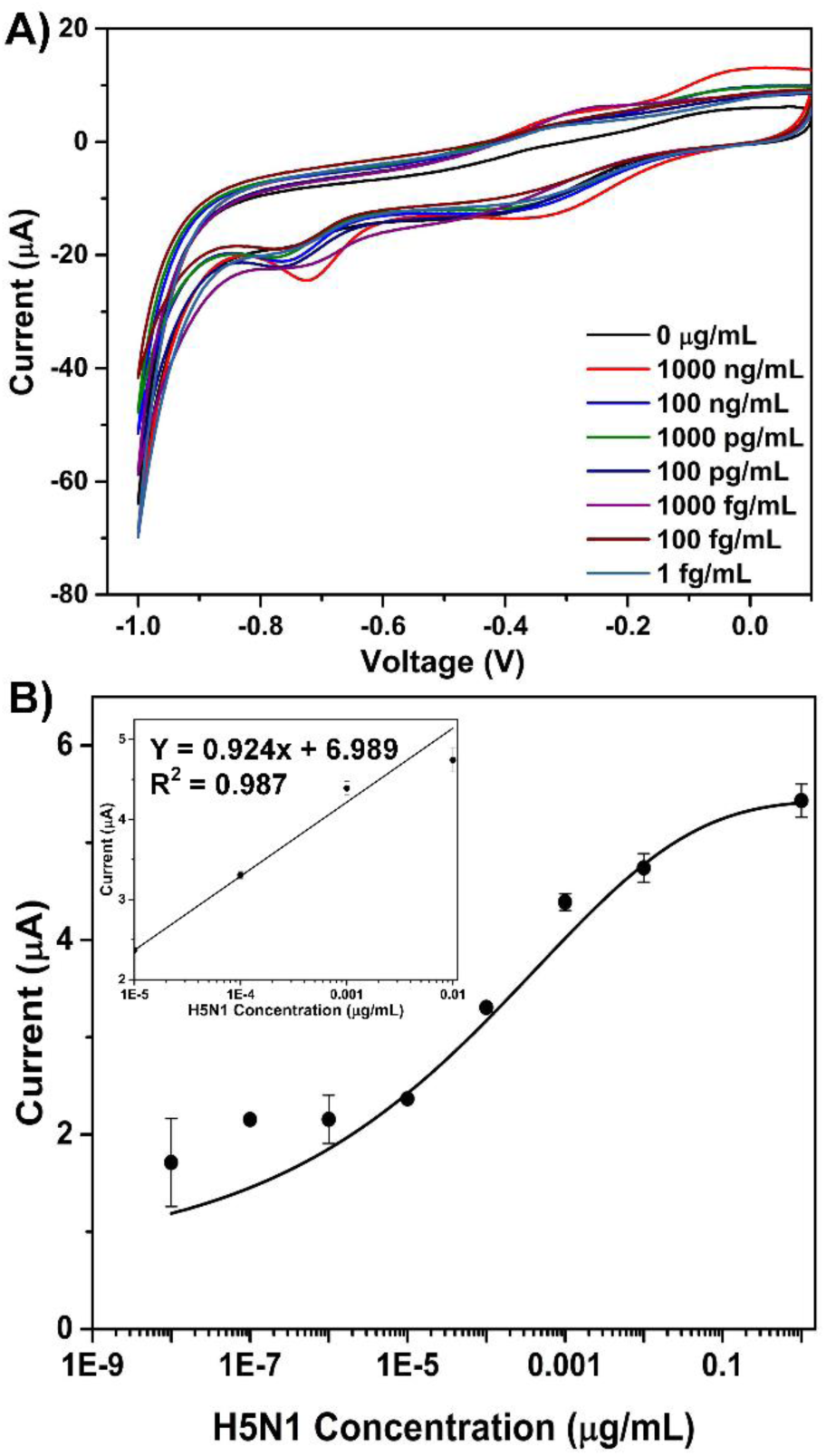
(A) Cyclic voltammetry profiles for H5N1 surface protein concentrations spiked in 1X PBS (pH 7.4) ranging from 0 µg/mL to 1 fg/mL. (B) Calibration curve of H5N1 surface protein immunosensing derived from the CV. Inset: Linear detection range from 10 ng/mL to 10 pg/mL.

### 3.4 H4N6 virus detection in blood

The goal of this study was to develop an immunosensor with the capability to detect avian influenza A (H4N6) virus in whole chicken blood. Upon conducting H4N6 sensing experiments in three-parts diluted whole chicken blood, it was found that the sensor can detect the viru It can be observed that a linear response exists for concentrations between 128 HAU and 1.28 × 10^−7^ HAU, with a sensitivity of 0.718 µA/HAU and a coefficient of determination of 0.975. Additionally, selectivity and cross sensitivity tests were conducted for H4N6 (Fig. 5).

These results highlight the selectivity of the immunosensor, as non-target viruses and viral proteins were statistically indistinguishable from a blank sample. More importantly, the immunosensor is selective to H4N6 even when exposed to H9N2 avian influenza virus. It can also be seen that there is no cross sensitivity with peptidoglycan, meaning that H4N6 can still be detected in the presence of peptidoglycan. These results suggest that the developed immunosensor is highly specific to H4N6 in whole chicken blood and that the sensor is ultrasensitive.

### 3.5 Validation with commercial ELISA kit

The sensitivity of the designed immunosensor was compared to those of a commercially available ELISA kit. The ELISA results for H4N6 inactivated protein demonstrate a sensitivity of 0.128 HAU (Fig. S6). Concentrations lower than 0.128 HAU are statistically indistinguishable by the ELISA kit. In comparison, the immunosensor (LOD = 1.28 × 10^−7^ HAU) was found to be 10^6^ times more sensitive than the commercial ELISA kit. Thus, the designed immunosensor exhibits ultra-sensitivity towards H4N6 inactivated virus.

## 4. Discussion

### 4.1 Nanocomposite characterization

From TEM images obtained (Fig. 2), it is evident that spikey/star shaped Au nanoparticles were formed. Due to one-pot in-situ nanocomposite synthesis, the Au nanoparticles bind to graphene electrostatically, thus reducing the overall synthesis time. These composites can provide enhanced effective surface area, superior catalytic properties, increased specificity, and limit of detection (LOD) in comparison to using graphene alone (Bai and Shen, 2012). As an example, individual graphene sheets tend to form irreversible clusters due to van der Waals forces and π-π stacking, thereby reducing their electrochemical properties (Stankovich et al., 2007). However, the incorporation of a second phase (i.e. non-spherical gold nanoparticles) provides a nano-spacer, which increases the graphene interlayer distance to minimize clumping. This effectively increases the conductance in two ways: the first being that both sides of graphene sheets are now accessible and the second being the addition of a conductive metal layer (Si and Samulski, 2008; Tien et al., 2010). The non-spherical confirmation will allow for increased surface area contacts between the nanoparticles and graphene sheet layers. Furthermore, nanospacing allows both sides of the graphene sheets to be conductive by reducing π – π stacking phenomena. The significance of the presented TEM images is that non-spherical Au nanoparticles were formed, and that the graphene sheets were well-decorated with nanoparticles; thus, forming non-spherical graphene – gold nanocomposites.

### 4.2 Antibody concentration optimization

The optimized antibody concentration that was used during testing was 1 µg/mL. However, as concentration was increased beyond 1 µg/mL, the associated peak current began to increase as well (Fig. S4). This contradicts electrochemical theory because as antibody concentration is increased, impedance of the electrode should also increase, thus reducing the peak current. These results may have been affected by steric hindrance as antibody concentration was increased. It may also be possible that the antibodies agglomerated together at higher concentrations, forming a conductive layer. Further studies are required to confirm whether these phenomena are occurring.

### 4.3 H5N1 surface protein detection in 1X PBS

The results obtained from the cyclic voltammetry studies with respect to recombinant protein concentration (Fig. 3(A)) exhibit baseline shifts, which make some higher concentration curves appear to have weaker peaks than some lower concentrations. This could be due to electrode to electrode variation resulting from non-uniform drying rate of nanocomposite. The resultant peak current values were used to develop a strong current – protein concentration relationship. The negative peaks at -0.75 V (Fig. 3) agree with previous work on the electrochemical reporting properties of CdTe (Amelia et al., 2012). The immunosensor possessed a lower LOD of 10 fg/mL and an upper LOD of detection of 1 µg/mL in PBS (pH 7.4) based on the obtained sigmoidal data (Fig. 3(B)). These results are agreeable with previously developed ELISA and nanoenzyme techniques (Ahmed et al., 2017). It is quite possible that the biosensor can detect concentrations greater than 1 µg/mL; however, higher concentrations have yet to be tested. It can be said that the linear concentration range (Fig. 3(B) inset) shows promise of ultra-sensitive detection of the target antigen.

### 4.4 H4N6 inactivated virus detection in blood

The immunosensor demonstrated ultra-sensitive detection of avian influenza A (H4N6) in three-parts diluted whole blood, with an upper and lower LOD of 128 HAU and 1.28 × 10^−7^ HAU (Fig. 4). The designed immunosensor also exhibited excellent selectivity towards H4N6 inactivated virus (Fig. 5). Even in the presence of highly concentrated peptidoglycan, a component of bacterial cell walls, H4N6 could be selectively detected. These results are highly significant because some samples (e.g. blood, faeces, mucosa, and sputum) can contain both viral and bacterial contaminants – it is important to be able to detect the target virus in such complex media. Due to the 10-minute incubation time, the time-to-results have been significantly reduced, when compared to conventional ELISA. Moreover, the immunosensor was 10^6^ times more sensitive than the commercial ELISA. To demonstrate the novelty of this sensor, it was also compared with previous works (Table 1).

**Fig. 4:**
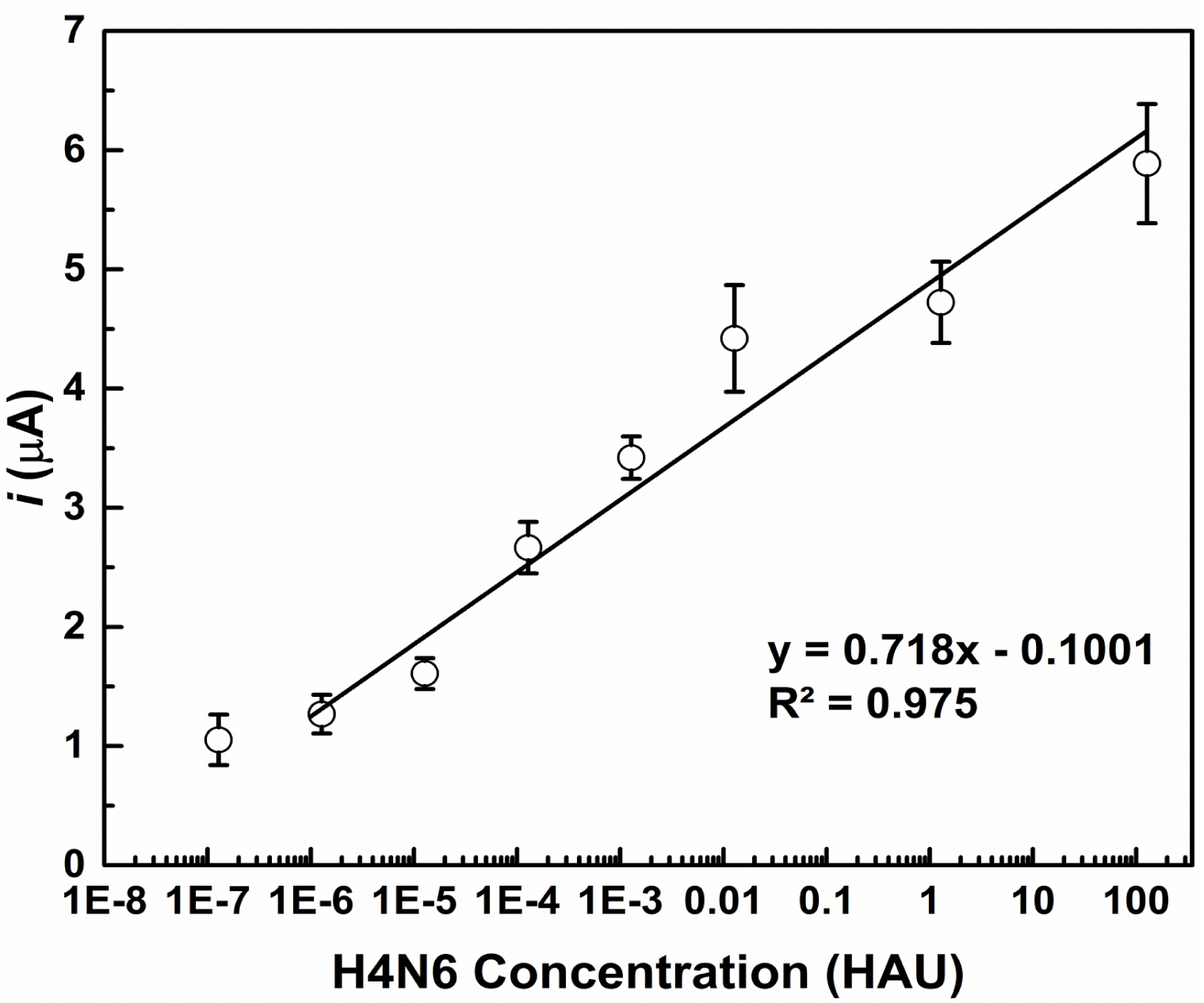
Avian influenza A (H4N6) sensing results in three-parts diluted whole chicken blood. The sensor exhibits a detection range from 128 HAU to 1.28 × 10^−7^ HAU in blood.

**Fig. 5:**
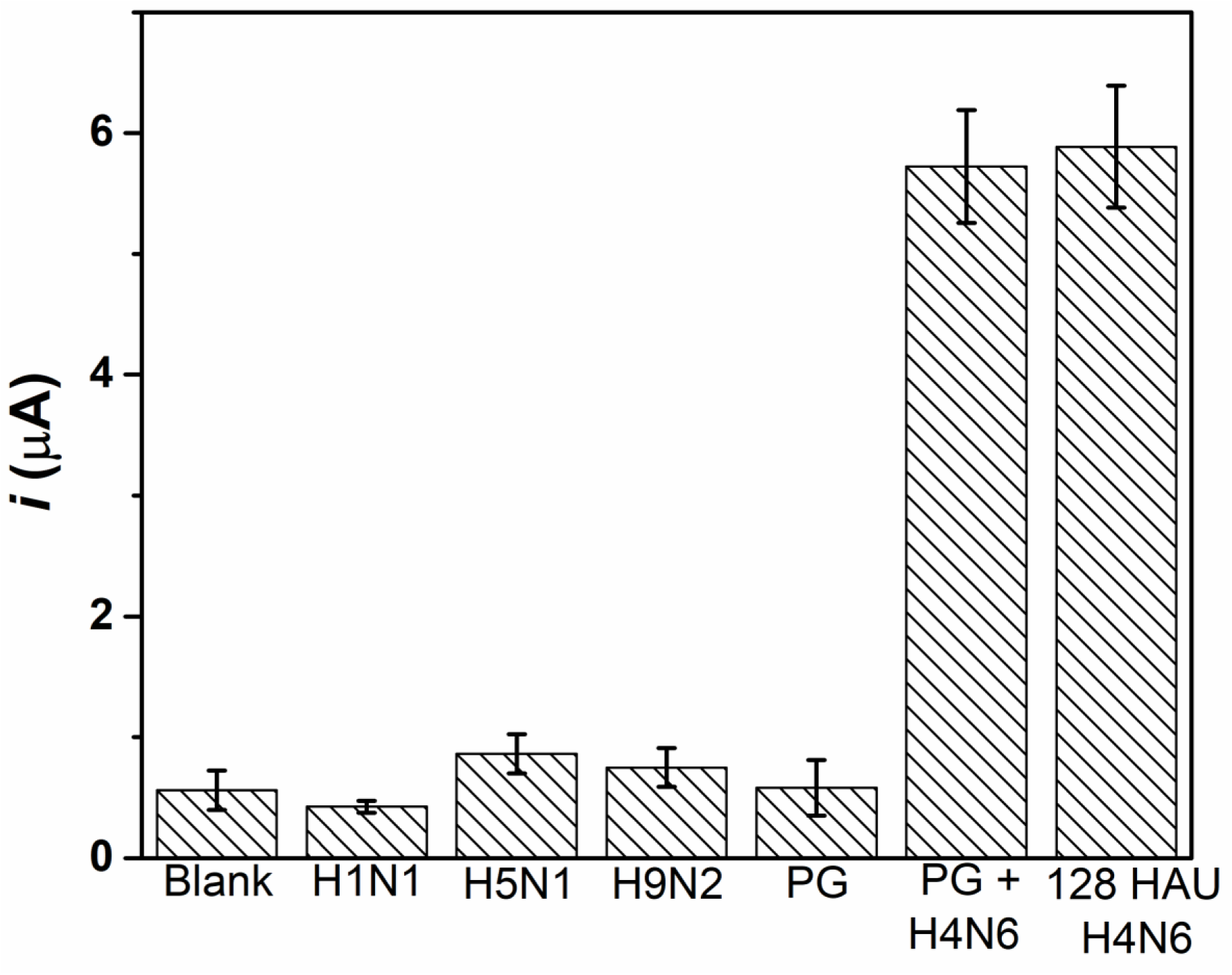
Selectivity and cross-sensitivity results for the developed avian influenza A (H4N6) virus immunosensing. The blank, H1N1 protein, H5N1 protein, peptidoglycan (PG), and H9N2 virus signals are statistically indistinguishable, thus, highlighting the selectivity of the immunosensor to the H4N6 antigen. Cross-sensitivity was observed between PG and H4N6 virus, however even in the presence of PG, H4N6 could be distinctly detected and measured.

**Table 1:** Comparison of this work to recent electrochemical-based influenza biosensor studies.

| Sensor (method) | Mechanism | Antigen(s) | Sample matrix | LOD | Detection range | Reference |
| --- | --- | --- | --- | --- | --- | --- |
| Electrochemical Immunosensor (CV, EIS, CA) | SPGE/RGO/CA/Ab/Ag on microfluidic chip | H1N1 virus | PBS | 0.5 PFU | 1 – 10<sup>4</sup> PFU | (Singh et al., 2017) |
| Electrochemical Immunosensor (N/A) | SPCE/Ab/BSA/Ag | H1N1 virus | chick embryo allantoic fluid | 0.43 HAU | 4 – 64 HAU | (Zhang et al., 2017) |
| Aptamer Sensor (CV, DPV) | SPCE/Au NP/DNA-aptamer/Ag/Ab-ALP | H5N1 protein | PBS | 100 fM | 100 fM – 10 pM | (Diba et al., 2015) |
| Electrochemical Immunosensor (DPV) | GCE/Au NP/Ab<sub>1</sub>/BSA/Ag/Ab<sub>2</sub>/Pt-pZnO-hemin | Not mentioned | PBS; 10 parts diluted human sera | 0.76 pg/mL | 0.001 – 60 ng/mL | (Yang et al., 2016) |
| Electrochemical Immunosensor (EIS, SWV) | SPGE/Au NP/scFV/BSA/Ag-His | H5N1 protein | PBS | 0.6 pg/mL | 4 – 20 pg/mL | (Góra-sochacka et al., 2016) |
| Electrochemical Immunosensor (CA) | Au electrode/ZnO NR/Ab<sub>1</sub>/BSA/Ag/Ab<sub>2</sub> on microfluidic chip | H1N1 protein; H5N1 protein; H7N9 protein | PBS | 1 pg/mL | 1 pg/mL – 10 ng/mL | (Han et al., 2016) |
| Electrochemical Immunosensor (CV, LSV) | Au electrode/ G-Au NP/Mab/Ag/PAb-Ag NP-G | H7 protein | PBS | 1.6 pg/mL | 1.6 pg/mL – 16 ng/mL | (Huang et al., 2016) |
| Electrochemical Immunosensor (CV) | SPGE/PL/G-Au NP-Ab/Ag/Ab-CdTe; SPCE/PL/G-Au NP-Ab/Ag/Ab-CdTe | H5N1 protein; H4N6 virus | PBS; 3 parts diluted chicken whole blood | 1 fg/mL; 1.28 x 10<sup>-7</sup> HAU | 1 fg/mL - 1 ug/mL; 1.28 x 10<sup>-7</sup> – 128 HAU | This work |

LOD – limit of detection, CV – cyclic voltammetry, EIS – electrochemical impedance spectroscopy, CA – chronoamperometry, DPV – differential pulse voltammetry, SWV – square wave voltammetry, LSV – linear sweep voltammetry, SPGE – screen-printed gold electrode, RGO – reduced graphene oxide, Ab – antibody, Ab<sub>1</sub> – primary antibody, Ab<sub>2</sub>- secondary antibody, Ag – antigen, SPCE – screen-printed carbon electrode, NP – nanoparticles, PL – L-polylysine, ALP – alkaline phosphatase, GCE – glassy carbon electrode, BSA – bovine serum albumin, pZnO – porous zinc oxide, scFV – single chain variable fragments, NR – nanorods, G – graphene, Mab – monoclonal antibody, PAb – polyclonal antibody, PBS – phosphate buffer solution, PFU – plaque forming units, HAU – hemagglutinating units.

This work is the first to detect real virus culture in a whole chicken blood sample matrix. Majority of the previously reported work first test the target analyte in buffer followed by spiking the target in biological fluid. This leads to inconsistency in results between buffered targets and spiked targets. Moreover, this work has demonstrated lower limits of detection than previously conducted studies. The goal of this design was to develop a sensing mechanism that is more sensitive than conventional ELISA and to reduce the time-to-results – both of which have been successfully accomplished in this study.

## 5. Conclusions

Due to the high virulence and zoonotic potential associated with H5 and H7 avian influenza pathotypes, it is of utmost importance to control outbreaks by reducing diagnostic turnaround. Current methods of detection exhibit many limitations with respect to sample handling/transport, expensive equipment and reagents, poor diagnostic turnaround and the need for specialized facilities. The work presented aims to provide a rapid electrochemical immunosensor that has potential to be used in a POC fashion on-site. The proposed immunosensor could be employed on farms in the form of a portable hand-held device. With such technology, farmers themselves could monitor the health of their flocks by simply taking a droplet of blood from their chickens, placing it onto a pre-coated electrode, and inserting the electrode into a reader.

Comparisons between conventional ELISA and recent electrochemical immunosensor studies were conducted. With respect to avian influenza A H5N1 recombinant protein, the imunosensor exhibited a lower LOD of 10 pg/mL and an upper LOD of 10 ng/mL in the linear range; however, a sigmoidal detection range from 10 pg/mL to 1 µg/mL was also established. With Respect to avian influenza A H4N6 virus, the immunosensor exhibited selectivity to H4N6, a lower LOD of 1.28 × 10^−7^ HAU, and an upper LOD of 128 HAU. In the case of H4N6 detection in blood, the immunosensor was found to be 10^6^ times more sensitive than its commercial ELISA counterpart. With respect to recent electrochemical immunosensor studies, this work is the first to perform total analysis (optimization and detection) of virus in a whole chicken blood matrix, which is a very complex media. The detection limits were much lower for this study in comparison to recent works. Thus, the designed immunosensor exhibited ultra-sensitivity in comparison to conventional ELISA methods and recent studies. In conclusion, this biosensor design is moving in the direction of rapid POC detection in blood, but this study is just the groundwork for a bigger and brighter future of avian influenza virus detection.

## Acknowledgments

The authors sincerely thank the Natural Sciences and Engineering Research Council of Canada (400705) and the Ontario Ministry of Agriculture, Food and Rural Affairs for funding this study (298634).

## References

Ahmed, S.R., Corredor, J.C., Nagy, É., Neethirajan, S., 2017. Amplified visual immunosensor integrated with nanozyme for ultrasensitive detection of avian influenza virus 1. doi:10.7150/ntno.20758

Amelia, M., Lincheneau, C., Credi, A., 2012. Chem Soc Rev Electrochemical properties of CdSe and CdTe quantum dots. R. Soc. Chem. 41, 5728–5743. doi:10.1039/c2cs35117j

American Plant Health Inspection Service, 2008. Avian Influenza Diagnostics and Testing Fact Sheet. https://www.aphis.usda.gov/publications/animal_health/content/printable_version/fsAI_diagnostics&testing.pdf

Bai, S., Shen, X., 2012. Graphene–inorganic nanocomposites. RSC Adv. 2, 64–98. doi:10.1039/c1ra00260k

Canadian Food Inspection Agency, 2015. Avian Influenza (AI) - What to expect if your animals are infected - Animals - Canadian Food Inspection Agency [WWW Document]. URL http://www.inspection.gc.ca/animals/terrestrial-animals/diseases/reportable/ai/what-to-expect-if-your-animals-are-infected/eng/1334853795705/1334853885674 (accessed 2.9.17).

Centers for Disease Control and Prevention, 2017. Avian Influenza A (H7N9) Virus | Avian Influenza (Flu) [WWW Document]. URL https://www.cdc.gov/flu/avianflu/h7n9-virus.htm (accessed 2.9.17).

Centers for Disease Control and Prevention, 2015. Highly Pathogenic Avian Influenza A (H5N1) in Birds and Other Animals | Avian Influenza (Flu) [WWW Document]. URL https://www.cdc.gov/flu/avianflu/h5n1-animals.htm (accessed 2.9.17).

Diba, F.S., Kim, S., Jin, H., 2015. Amperometric bioaffinity sensing platform for avian influenza virus proteins with aptamer modified gold nanoparticles on carbon chips 72, 355–361. doi:10.1016/j.bios.2015.05.020

Góra-sochacka, A., Sirko, A., Dehaen, W., Jarocka, U., Radecki, J., Radecka, H., 2016. An electrochemical immunosensor based on a 4, 4-thiobisbenzenethiol self-assembled monolayer for the detection of hemagglutinin from avian influenza virus H5N1 228, 25–30. doi:10.1016/j.snb.2016.01.001

Han, J., Lee, D., Hong, C., Chew, C., Kim, T., Jungho, J., 2016. A multi-virus detectable microfluidic electrochemical immunosensor for simultaneous detection of H1N1, H5N1, and H7N9 virus using ZnO nanorods for sensitivity enhancement. Sensors Actuators B. Chem. 228, 36–42. doi:10.1016/j.snb.2015.07.068

Health canada, 2008. Avian Influenza (Bird Flu) Fact Sheet.

Huang, J., Xie, Z., Xie, Z., Luo, S., Xie, L., Huang, L., Fan, Q., Zhang, Y., Wang, S., Zeng, T., 2016. Silver nanoparticles coated graphene electrochemical sensor for the ultrasensitive analysis of avian influenza virus H7. Anal. Chim. Acta 913, 121–127. doi:10.1016/j.aca.2016.01.050

Jensen, T.H., Ajjouri, G., Handberg, K.J., Slomka, M.J., Coward, V.J., Cherbonnel, M., Jestin, V., Lind, P., Jorgensen, P.H., 2013. An enzyme-linked immunosorbent assay for detection of avian influenza virus subtypes H5 and H7 antibodies. Acta Vet Scand 55, 84. doi:10.1186/1751-0147-55-84

Neethirajan, S., Ahmed, S.R., Chand, R., Buozis, J., Nagy, É., 2017. Recent Advances in Biosensor Development for Foodborne Virus Detection 1. doi:10.7150/ntno.20301

Olsen, B., Munster, V.J., Wallensten, A., Waldenström, J., Osterhaus, A.D.M.E., Fouchier, R.A.M., 2006. Global patterns of influenza a virus in wild birds. Science 312, 384–8. doi:10.1126/science.1122438

Si, Y., Samulski, E.T., 2008. Exfoliated graphene separated by platinum nanoparticles. Chem. Mater. 20, 6792–6797. doi:10.1021/cm801356a

Singh, R., Hong, S., Jang, J., 2017. Label-free Detection of Influenza Viruses using a Reduced Graphene Oxide-based Electrochemical Immunosensor Integrated with a Microfluidic Platform. Nat. Publ. Gr. 2017, 1–11. doi:10.1038/srep42771

Singh, S., Alkie, T., Abdelaziz, K., Hodgins, D., Novy, A., Nagy, É., Sharif, S., 2016. Characterization of Immune Responses to an Inactivated Avian Influenza Virus Vaccine. Viral Immunol. 29, 269–275. doi:10.1089/vim.2015.0144

Stankovich, S., Dikin, D.A., Piner, R.D., Kohlhaas, K.A., Kleinhammes, A., Jia, Y., Wu, Y., Nguyen, S.T., Ruoff, R.S., 2007. Synthesis of graphene-based nanosheets via chemical reduction of exfoliated graphite oxide. Carbon N. Y. 45, 1558–1565. doi:10.1016/j.carbon.2007.02.034

Szretter, K.J., Balish, A.L., Katz, J.M., 2006. Influenza: propagation, quantification, and storage. Curr Protoc Microbiol Chapter 15, Unit 15G 1. doi:10.1002/0471729256.mc15g01s3

Tien, H., Huang, Y., Yang, S., Wang, J., Ma, C.M., 2010. The production of graphene nanosheets decorated with silver nanoparticles for use in transparent, conductive films. Carbon N. Y. 49, 1550–1560. doi:10.1016/j.carbon.2010.12.022

United States Department of Agriculture, 2015. Avian Influenza Testing and Diagnostics Fact Sheet [WWW Document]. USDA Press Off. URL https://www.usda.gov/documents/usda-avian-influenza-diagnostics-testing-factsheet.pdf (accessed 2.9.17).

Veerapandian, M., Hunter, R., Neethirajan, S., 2016a. Lipoxygenase-modified Ru-bpy / graphene oxide: Electrochemical biosensor for on-farm monitoring of non-esterified fatty acid. Biosens. Bioelectron. 78, 253–258. doi:10.1016/j.bios.2015.11.058

Veerapandian, M., Hunter, R., Neethirajan, S., 2016b. Dual immunosensor based on methylene blue-electroadsorbed graphene oxide for rapid detection of the influenza A virus antigen. Talanta 155, 250–257. doi:10.1016/j.talanta.2016.04.047

Veerapandian, M., Neethirajan, S., 2015. Graphene oxide chemically decorated with Ag–Ru/ chitosan nanoparticles: fabrication, electrode processing and immunosensing properties. RSC Adv. 5, 75015–75024. doi:10.1039/C5RA15329H

Weng, X., Neethirajan, S., 2016. A micro fluidic biosensor using graphene oxide and aptamer- functionalized quantum dots for peanut allergen detection. Biosens. Bioelectron. 85, 649–656. doi:10.1016/j.bios.2016.05.072

World Health Organization, 2017a. WHO | Influenza (Seasonal) [WWW Document]. WHO. URL http://www.who.int/mediacentre/factsheets/fs211/en/ (accessed 3.9.17).

World Health Organization, 2017b. WHO | Avian and other zoonotic influenza.

World Organization for Animal Health, 2016. Avian Influenza (Infection with Avian Influenza Viruses), in: Manual of Diagnostic Tests and Vaccines for Terrestrial Animals. OIE, pp. 1–23.

Yang, Z., Zhuo, Y., Yuan, R., Chai, Y., 2016. A nanohybrid of platinum nanoparticles-porous ZnO – hemin with electrocatalytic activity to construct an amplified immunosensor for detection of influenza. Biosens. Bioelectron. 78, 321–327. doi:10.1016/j.bios.2015.10.073

Zhang, H., Zhang, S., Liu, N., 2017. Prevention and control of emergent infectious disease with high specific antigen sensor. Artif. Cells, Nanomedicine, Biotechnol. 0, 1298–1303. doi:10.3109/21691401.2016.1161638

Zhu, L., Zhu, C., Deng, G., Zhang, L., Zhao, S., Lin, J., Li, L., Jiao, P., Liao, M., Liu, Y., 2014. Rapid identification of H5 avian influenza virus in chicken throat swab specimens using microfluidic real-time RT-PCR. Anal. Methods 6, 2628. doi:10.1039/c3ay42126k

